# Integration of Multi-omics Data for the Classification of Glioma Types and Identification of Novel Biomarkers

**DOI:** 10.1101/2023.12.22.572983

**Authors:** Francisca G. Vieira, Regina Bispo, Marta B. Lopes

## Abstract

Glioma is currently one of the most prevalent types of primary brain cancer. Given its high level of heterogeneity along with the complex biological molecular markers, many efforts have been made to accurately classify the type of glioma in each patient, which, in turn, is critical to improve early diagnosis and increase survival. Nonetheless, as a result of the fast-growing technological advances in high throughput sequencing and evolving molecular understanding of glioma biology, its classification has been recently subject to significant alterations. In this study, we integrate multiple glioma omics modalities (including mRNA, DNA methylation, and miRNA) from The Cancer Genome Atlas (TCGA), while using the revised glioma reclassified labels, with a supervised method based on sparse canonical correlation analysis (DIABLO) to discriminate between glioma types. We were able to find a set of highly correlated features distinguishing glioblastoma from lower-grade gliomas (LGG) that were mainly associated with the disruption of receptor tyrosine kinases signaling pathways and extracellular matrix organization and remodeling. On the other hand, the discrimination of the LGG types was characterized primarily by features involved in ubiquitination and DNA transcription processes. Furthermore, we could identify several novel glioma biomarkers likely helpful in both diagnosis and prognosis of the patients, including the genes *PPP1R8, GPBP1L1, KIAA1614, C14orf23, CCDC77, BVES, EXD3, CD300A* and *HEPN1*. Overall, this classification method allowed to discriminate the different TCGA glioma patients with very high performance, while seeking for common information across multiple data types, ultimately enabling the understanding of essential mechanisms driving glioma heterogeneity and unveiling potential therapeutic targets.

## 1 Introduction

Glioma is one of the most common malignant primary brain tumors in adults, comprising multiple types with distinct biomolecular characteristics. Given the low overall patient survival, there is an urgent growing need for new effective treatment approaches. However, the marked differences in prognosis and therapeutic outcomes still represent major clinical challenges [1]. To understand the heterogeneity among gliomas, tremendous efforts have been made to identify different types [2, 3, 4, 5]. Although the tumors have been historically designated according to morphological and histopathological traits, the World Health Organization (WHO) Classification of Tumors of the Central Nervous System (CNS) has already started to incorporate additional driver molecular markers in their revised guidelines, obtained via information extracted from recent sequencing technologies, such as gene expression and DNA methylome profiling, to further refine their characterization [6]. Adult-type diffuse gliomas are now broadly classified based on their molecular profile: isocitrate dehydrogenase (IDH)-mutant samples can be labeled as either astrocytoma (grades 2, 3 and 4) or oligodendroglioma (grades 2 and 3) based on whether there is 1p/19q codeletion or not, respectively, while the IDH-wildtype with certain genetic parameters (e.g. *TERT* promoter mutation or *EGFR* gene amplification) are classified as glioblastoma (grade 4) [7, 8]. Tumors may also be referred to as Lower-Grade Gliomas (LGG) according to a lower proliferative activity (grades 1 to 3), with the exception of glioblastoma (GBM) which exhibits high malignancy traits.

Recent advances in high-throughput technologies, particularly next-generation sequencing, have been allowing the extraction of an increasing quantity of biological data, which is not limited to one but instead includes multiple omics modalities collected from the same individuals. The joint integrative study of the different but linked layers of genetic regulation (e.g., genome, epigenome, transcriptome, proteome, metabolome) offers an opportunity to build a more comprehensive landscape of biological systems and further enhance the molecular understanding of disease, where research has relied mostly on single-omics data [9].

Several methods have been proposed to address multi-omics data integration and analysis. Joint dimensionality reduction (JDR) techniques transform the data into a lower-dimensional space based on matrix decomposition, learning latent factors that capture biological sources of common variability across various omics datasets. These techniques include, for instance, sparse variants of both Canonical Correlation Analysis (CCA) [10, 11] and Multi-block Partial Least Squares (PLS) [12], joint Non-negative Matrix Factorization (NMF) [13], Joint and Individual Variation Explained (JIVE) [14, 15] and Multiomics Factor Analysis (MOFA) [16, 17]. Furthermore, supervised extensions of CCA have been recently suggested in a classification framework to identify relevant multi-view features that are not only highly correlated but also optimally separate subjects into distinct groups [18, 19, 20]. In particular, Data Integration Analysis for Biomarker discovery using Latent cOmponents (DIABLO) [18] has been used in multiple omics applications, including the identification of disease mechanisms and biomarkers [21, 22]. Moreover, a recent benchmark analysis in cancer type classification [23] revealed that DIABLO performed significantly better than several other methods, including MOFA/MOFA+ [17], principal component analysis (PCA) and iClusterBayes/iClusterPlus/iCluster [24].

In this work, an integrative multi-layer omics study using mRNA, DNA methylation, and miRNA data from The Cancer Genome Atlas (TCGA) was performed to classify and characterize the different glioma types, considering its recently updated reclassification [7, 8]. Firstly, in Section 2, we describe the data collected and detail all the methods used in this work. We used DIABLO to select a subset of highly correlated features from the different modalities relevant to distinguish either the three glioma types (as present in Section 3.1) or specifically to discriminate the two LGG types (Section 3.2). This is particularly relevant considering the ongoing evolution in glioma classification and the need of identifying novel genetic markers that aid in its characterization. Aiming to delineate unknown mechanisms of glioma pathobiology and unveil novel therapeutic targets, we further investigated the correlation between the selected features, the pathways involved, and the prognostic value based on their effect in the survival probability of the patients (Section 3.3). Lastly, in Sections 4 and 5, we discuss the obtained results and highlight the main conclusions of the work developed, respectively.

## 2 Materials and Methods

### 2.1 Data Description

In this study, we used the GBM and LGG datasets from TCGA, available in the Genomic Data Commons Data Portal with project names TCGA-GBM and TCGA-LGG. The TCGA level-3 data on gene expression (Illumina mRNAseq), DNA methylation (Illumina HumanMethylation450), miRNA expression (Illumina miRNAseq, BCGSC miRNA Profiling workflow) and clinical information were retrieved. RNA-seq data normalized for gene length and for sequencing depth (Transcript per million, TPM) with upper quantile normalization was extracted for DIABLO and survival analysis. RNA-seq RSEM expected counts, without normalization, were extracted for differential gene expression analysis. The R packages RTCGAToolbox v2.28.1 [25] and TCGAbiolinks 2.25.3 [26] were used to collect the data.

DNA methylation data was first preprocessed by removing non-valid entries (start = end = -1), probes mapping multiple places, non-CpG sites, sexual chromosome and missing (NA) entries. For each dataset, features with zero variance or having very few unique values relative to the number of samples (cutoff of 10%) and a large ratio of the frequency of the most common value to the frequency of the second most common value (cutoff ratio of 95/5) were filtered. The intersection of the datasets was done to keep only the samples that were present in all the datasets. From the 278 GBM individuals, 108 had both mRNA and DNA methylation data, and 5 had miRNA data, so the latter data type was not used in GBM discrimination. From the 262 astrocytoma individuals, 248 had both mRNA and DNA methylation data, and 243 had miRNA data, while from the 169 oligodendroglioma individuals, 166 had mRNA, DNA methylation and miRNA data, characterized respectively by 19068, 297602 and 1118 filtered features.

In the classification task, the data was split into train and test subsets, in a predefined ratio (70% for training and 30% for testing), while preserving relative ratios of the different classes (the different glioma types are in the same proportion as the original dataset).

### 2.2 Data Integration Analysis for Biomarker discovery using Latent cOmponents - DIABLO

Data Integration Analysis for Biomarker discovery using Latent cOmponents (DIABLO) [18] was applied in this work, using the R package mixOmics v6.22.0 [27], to select co-expressed variables from the different datasets that discriminate between the glioma types. DIABLO discriminant analysis extends the multivariate methodology Generalized Canonical Correlation Analysis (GCCA) [11] to a supervised framework by replacing one data matrix *X* ^(*q*)^ with the outcome dummy matrix *Y*. For *Q* omics datasets measuring the expression levels of the P_q_ omics variables on the same biological samples, GCCA solves for each component (canonical variate) *h*:

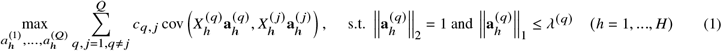

where 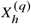 and 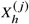 represent the datasets, 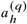 and 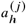 are the loading vectors (component coefficients) on component *h* with *q, j* = 1,…,*Q* (*q* ≠ *j*). For all **a** ∈ ℝ^*n*^, *ℓ*_1_ and *ℓ*_2_ norms are, respectively, defined by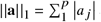 and 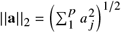. *λ* ^(*q*)^ is a *ℓ*_1_ penalisation parameter to induce sparsity on the loading vector **a**_*h*_ by shrinking some coefficients to zero. One data matrix *X* ^(*q*)^ is given by the outcome dummy matrix, *Y*, indicating the class membership of each individual. *C* = {*c*_*q, j*_ }_*q, j*_ is the design matrix, a Q×Q matrix that specifies whether datasets should be correlated and includes values between zero (datasets are not connected) and one (datasets are fully connected), enabling to model a particular relation between pairs of omics data and to constraint the model to only take into account those specific pairwise covariances, as expected from prior biological knowledge or experimental design. The design matrix was specified based on a data-driven approach. Using PLS as a preliminary analysis, integrating two datasets at a time to assess the common information between them, the correlation value was then used in *c*_*q,j*_ for *q, j* < *Q*, while *c*_*q,j*_ = 1 for *q, j* = *Q*.

In this work, 2 components were used, since in general, *K - 1* components are sufficient to achieve the best classification performance, where *K* is the number of classes [27]. It was also verified, by evaluating the difference in overall misclassification error rate, that there was no gain in performance when adding more components to the sparse model.

When using sparse GCCA, the component scores *t*_*h*_ = *X*_*h*_*a*_*h*_ are defined on a small subset of variables with non-zero coefficients, leading to variable selection that aims to optimally maximize the discrimination between the *K* outcome classes in *Y*. In DIABLO, a soft-thresholding is used, replacing the non negative parameter *λ*^(*q*)^ that controls the amount of shrinkage in *a*_*h*_ by the number of features to select on each dimension [27]. To select the number of features, a step-by-step approach was used, by assessing the performance of the model (measured via overall misclassification error rate) for each value of features provided as a grid, one component at a time, using 5x5 cross-validation (CV). First, a grid with a higher amplitude of values but low resolution was evaluated, and subsequently finer grids were analyzed around the previously selected values with progressively smaller steps, until the values converged.

Considering an independent test set, the predicted coordinates (scores) are computed for each new observation on the set of H latent components, and then used to predict each of the dummy variables. The final predicted class is then obtained, minimizing the distance to the centroid. The predictions are combined by weighted vote, where each omics dataset weight is defined as the correlation between the latent components associated to that particular data set and the outcome, from the training set. The final prediction is the class that obtains the highest weight across all datasets.

In this work, 30 different partitions of the data were considered, retrieving from each trained model the variables with non-zero coefficients and their absolute values, such as the performance measures (accuracy, precision, recall, F1) after predicting the labels in each test set. The receiver operating characteristic (ROC) curves were also used to evaluate the performance of each model and each component in predicting the glioma type, and the respective area under the curve (AUC) was determined.

### 2.3 Selection of Features and Database Search

For further analyzes we have selected the variables that, within the 30 different models, occurred more frequently in the components with non-zero coefficient and with higher median absolute value: frequency *>* 15 and median absolute value *>* 0.05 or frequency *>* 10, and median absolute value *>* 0.2. The outcome class of each feature (the glioma type for which the expression of that feature is more distinguished from the other types) was determined as follows: For the RNA features, the respective outcome type was determined by differential gene expression analysis, where, for each gene, the outcome is the class in which it is more differentially expressed (with lower false discovery rate (FDR) and higher log(FC)). For the methylation features, the outcome was determined for each methylation site as the class with highest averaged methylation level difference to the other two classes. For the miRNA features, the outcome was determined as the class with higher averaged miRNA expression (in this case there were only the two LGG types).

Each selected variable was searched for associations with glioma and cancer over different databases. In the case of DNA methylation features, they were first mapped back to gene symbols, using methylGSA v1.16.0 package [28] and the multi-symbol checker tool from HUGO Gene Nomenclature Committee. The search was performed using all the human collections from the Molecular Signatures Database (MolSigDB) [29] and OMIM^®^ [30] with the keywords “glioma”, “glioblastoma”, “oligodendroglioma”, “astrocytoma” and in PubMed^®^, using as keywords the symbol of each gene and “AND glioma” or “AND cancer”. The gene targets of each miRNA were determined using the MiRTarBase database [31].

### 2.4 Differential Expression Analysis

Differential expression analysis was performed using the edgeR package v3.40.2 [32]. Genes with very low counts were first filtered, and trimmed mean of M values (TMM) normalization was applied to account for compositional biases [33]. Differential expression between the experimental groups was tested using quasi-likelihood F-test (considering a negative binomial distribution). Genes with an FDR-adjusted p-value (based on Benjamini and Hochberg (BH) adjustment [34]) of less than 0.05 were deemed significantly differentially expressed genes (DEG). Furthermore, to define a compromise between statistical significance and biological variation, the genes for which:

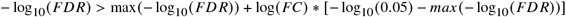

were considered positively differentially expressed while the genes for which:

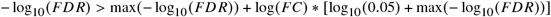

were considered negatively differentially expressed. All the remaining DEGs, although significant were considered not relevant given the low log(FC).

### 2.5 Overrepresentation and Gene Set Enrichment Analysis

Differentially expressed genes that demonstrated significant fold changes between conditions were screened for KEGG pathways and Gene Ontology (GO) terms with over-representation enrichment analysis (ORA) in the software WebGestalT [35]. Using the same software, the full list of expressed genes ranked by a combination of statistical significance and fold-change:

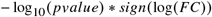

was used in gene set enrichment analysis (GSEA). Terms and pathways were considered significant for a FDR < 0.05 (BH adjustment).

### 2.6 Correlation and Survival Analysis

The Pearson correlation between each pair of selected features was computed using the respective values of expression in all the patients, with the R package Hmisc v5.0-1 [36] and plotted using the R package pheatmap v1.0.12 [37].

A Kaplan-Meier survival curve was estimated using the clinical data from the patients of the TCGA-LGG and TCGA-GBM projects, with the R packages survival v3.5-3 for the analysis and survminer v0.4.9 for plotting [38, 39]. For each evaluated gene (RNA and DNA methylation selected features), the tumor samples were split into two groups: low and high expression levels, using either the median value as cut-off or when existing two distinct density peaks, the threshold is set as the expression for which the density has the local minimum between the peaks. The statistical significance of survival differences in the analysis was assessed using the log-rank test, using the 0.05 significance level.

## 3 Results

### 3.1 Integration of two omics layers for the classification of glioma types

To discriminate between the three glioma types, namely GBM, astrocytoma, and oligodendroglioma, two components of mRNA and DNA methylation features were first obtained using DIABLO.

Regression analysis with PLS showed that the datasets were highly correlated (cor = 0.8796), and therefore the design matrix was set with a weight of 0.8. Based on the grid search, the number of mRNA variables with non-zero coefficient that was chosen were 41 (out of 19068) for the first component and 7 for the second component, while the number of DNA methylation variables chosen was 11 (out of 297602) for the first component and 41 for the second component.

The performance metrics evaluating the classification of the glioma types, based on the obtained components, are summarized in Table 1. Keeping only two components was sufficient to achieve a good overall accuracy of 98%, with the first component primarily differentiating GBM from the LGG and the second component mostly distinguishing between astrocytoma and oligodendroglioma, as depicted by Figure 1A. It is clear that GBM is considerably easier to identify since it has the highest F1 value and its ROC curve separating from the LGG types has an area under the curve close to 1 in all the components (Figure 1B).

**Table 1:** Performance metrics for the classification of the three glioma types using two components based on RNA and DNA methylation features.

| Glioma type | Performance metrics |  |  |  |
| --- | --- | --- | --- | --- |
|  | Precision | Recall | F1 | Accuracy |
| Astrocytoma | 0.996 | 0.964 | 0.980 | 0.981 |
| Glioblastoma | 0.995 | 0.992 | 0.993 |  |
| Oligodendroglioma | 0.952 | 1.000 | 0.976 |  |

**Figure 1:**
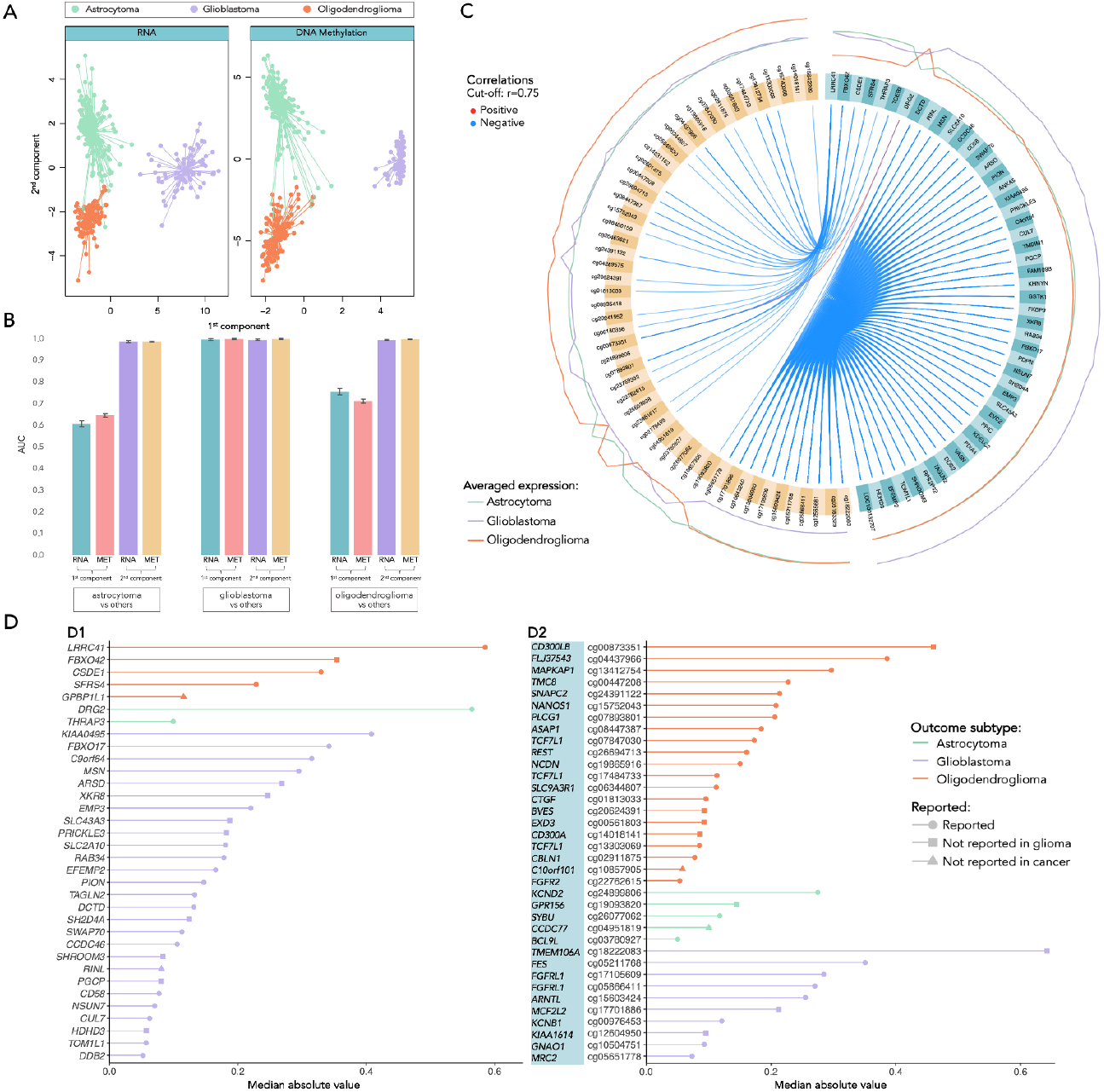
Separation and classification of three glioma types based on components with selected mRNA and DNA methylation features and respective performance. A: Separation of the individuals based on the first two components (orange dots represent the oligodendroglioma, green dots represent the astrocytoma and lilac dots represent the GBM individuals). B: AUC of the ROC curves to discriminate each class vs the two others. C: Circos plot representing the averaged expression of the genes in each glioma type and correlations between the features selected in each of the data blocks (mRNA features colored in turquoise, and DNA methylation features in yellow). D: Features (D1: mRNA; D2: DNA methylation) selected by frequency and absolute median value, categorized by the outcome glioma type (oligodendroglioma colored in orange, astrocytoma in green and GBM in lilac). The symbol at the end of each line indicates: if already reported in glioma studies (circle); if not reported in glioma but in other types of cancer (square); if never reported in cancer at all (triangle), based on search on Human MSigDB Collections, OMIM, and PubMed.

Figure 1C shows the averaged expression levels of the selected features generally distinguish one of the classes from the other two (denoted as the outcome type). Moreover, features are highly correlated between blocks, where the methylation of some sites is mainly negatively correlated with the expression of other genes. The highly correlated blocks are displayed in Figure 3C and will be further described in Section 2.3.

We have then selected the variables that more frequently occurred in the components with non-zero coefficients and with higher loading. The selected variables are shown in Figure 1D (mRNA features in Fig.1D1 and DNA methylation features in Fig.1D2), ordered by the respective importance in the components (median absolute value) and grouped by outcome. The barplots also show whether each feature was reported in the literature or in biological databases with connections with glioma or cancer. Indeed, nearly all have been reported in cancer studies, and the vast majority have also been investigated in glioma. Moreover, most of the gene features selected were differentially expressed in GBM, while the selected methylation sites were more relevant for the discrimination of oligodendroglioma. For instance, some of the features that revealed to have the greatest influence in the distinction of GBM were the genes *KIAA0495, FBXO17, C9orf64, MSN* and *ARSD* and the methylated sites cg18222083 (*TMEM106A*), cg05211768(*FES*), cg17105609 and cg05866411 (*FGFRL1*), and cg15603424 (*ARNTL*), all reported in glioma studies with the exception of the genes *ARSD* and *TMEM106A*, which were only reported in other types of cancer. On the other hand, to discriminate astrocytoma, much fewer features were found to be relevant, including only two genes *DRG2* and *THRAP3*, both already reported in glioma studies, and the methylation sites cg24899806 (*KCND2*), cg19093820 (*GPR156*), cg26077062 (*SYBU*), cg04951819 (*CCDC77*) and cg03780927 (*BCL9L*), where *GPR156* and *CCDC77* were never reported in glioma, and the latter also never reported in any other cancer studies. Interestingly, two of the most important features to distinguish oligodendroglioma, the gene *FBX042* and the methylated gene *CD300LB* were also never reported in glioma studies.

### 3.2 Discrimination of Lower-grade glioma types using three omics layers

Once shown in the previous analysis that the LGG were more challenging to distinguish, a further analysis keeping only these two classes and including an additional data view - miRNA - was performed with DIABLO, again using only two components.

Regression analysis with PLS showed smaller correlation between DNA methylation and miRNA (cor = 0.6027) than the one obtained for the other pairs of data, and therefore the design matrix was set with a weight of 0.6 for this dataset pair and 0.8 for the others. Based on the grid search, the number of mRNA variables with non-zero coefficient that was chosen was 7 in both components, 29 DNA methylation variables for the first component and 42 for the second component, and 11 miRNA variables in both components.

The performance metrics evaluating the classification of the glioma types, based on the obtained components, are present in Table 2 and the AUC obtained are shown in Table 3. An accuracy of 97% was achieved, and the first component was able to distinguish by itself the classes (Figure 2A). Indeed, when observing the circos plot in Figure 2B, we can also see that the averaged expression levels of the selected features do not always separate the two classes (the lines of expression of the second component features tend always to overlap). The features are highly correlated between blocks, where the DNA methylation features are mainly negatively correlated with the expression of the genes, while there are many positive correlations between miRNA and both genes and methylation features.

**Figure 2:**
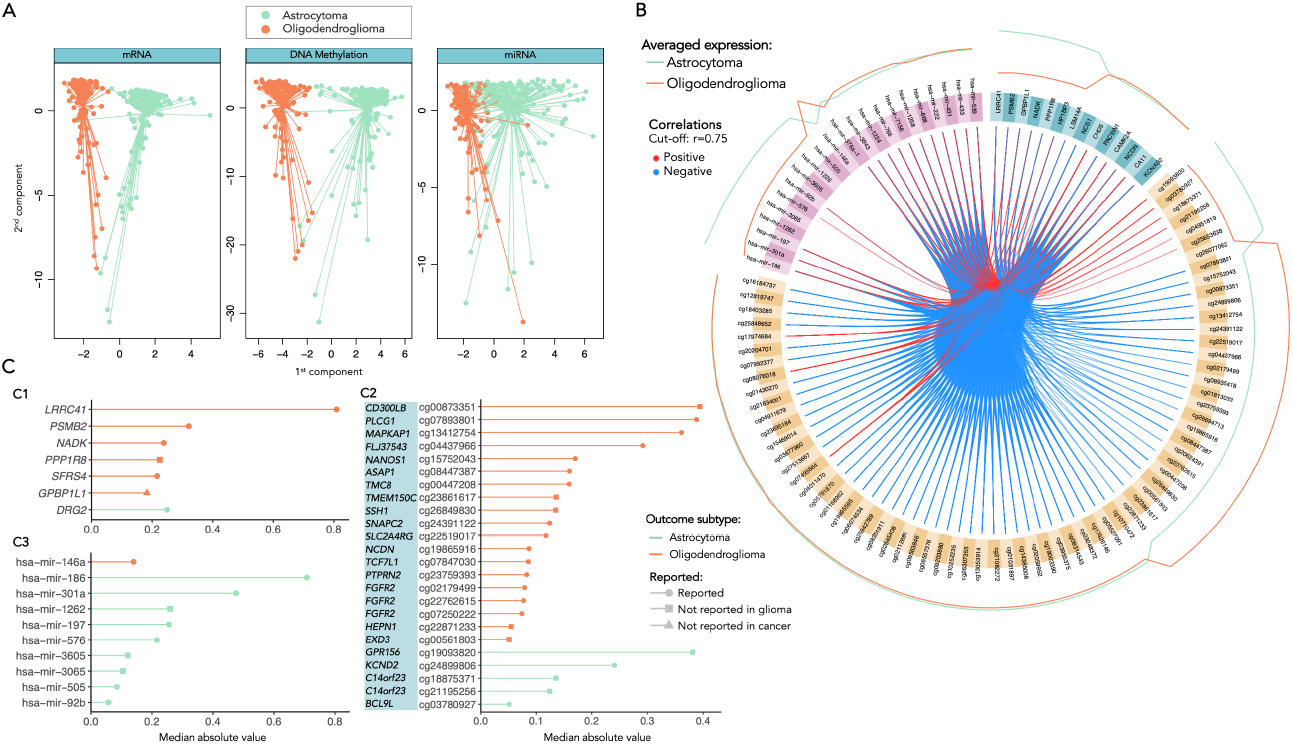
Separation and classification of LGG glioma types based on components with selected mRNA, DNA methylation and miRNA features. A: Separation of the individuals based on the first two components (orange dots represent the oligodendroglioma individuals, and green dots represent the astrocytoma individuals). B: Circos plot representing the averaged expression of the genes in each glioma type (orange line for oligodendroglioma and green line for astrocytoma) and correlations between the features selected in each of the data blocks (mRNA features colored in turquoise, DNA methylation features in yellow and miRNA features in pink). C: Features (C1: mRNA; C2: DNA methylation; C3: miRNA) selected by frequency and absolute median value, categorized by the outcome glioma type (oligodendroglioma colored in orange and astrocytoma in green). The symbol at the end of each line indicates: if already reported in glioma studies (circle); if not reported in glioma but in other types of cancer (square); if never reported in cancer at all (triangle), based on search on Human MSigDB Collections, OMIM, and PubMed.

**Table 2:** Performance metrics for the classification of the LGG types using two components based on mRNA, DNA methylation and miRNA features.

| Glioma type | Performance metrics |  |  |  |
| --- | --- | --- | --- | --- |
|  | Precision | Recall | F1 | Accuracy |
| Astrocytoma | 0.996 | 0.958 | 0.977 | 0.973 |
| Oligodendroglioma | 0.943 | 0.995 | 0.968 |  |

**Table 3:** AUC obtained for each component and each data layer for the classification of LGG types.

|  | AUC |  |  |
| --- | --- | --- | --- |
|  | mRNA | DNA methylation | miRNA |
| Component 1 | 0.994 | 0.987 | 0.995 |
| Component 2 | 0.994 | 0.931 | 0.887 |

We have then selected only the variables that occurred more frequently in the first component with non-zero coefficient, given that the ones selected by the second component were not useful to distinguish the classes. The selected variables are shown in Figure 2C (mRNA features in Fig.2C1, DNA methylation features in Fig.2C2 and miRNA features in Fig.2C3), ordered by the respective importance in the components (median absolute value) and grouped by outcome. Note that the selected features are not exactly the ones already selected before when only using mRNA and DNA methylation data (shown in the previous section 2.1). Indeed, the selection still includes some genes that were already chosen before: *LRRC41, SFRS4, GPBP1L1* and *DRG2*, highlighting their relevance for the separation of the classes, while also adding the genes *PSMB2, NADK* and *PPP1R8*, but now discarding *FBX042* and *CSDE1* as relevant genes. Also in the methylation sites, the more relevant methylated genes, such as *CD300LB, PLCG1, MAPKAP1, FLJ37543, NANOS1, ASAP1, TMC8* and *SNAPC2* are maintained in the selection, while more differences are detected within the comparison of features with lower absolute value. Furthermore, most of the genes and methylation sites are associated with the distinction of the oligodendroglioma class, while the majority of the selected miRNA are more expressed in astrocytoma. It is also interesting to note that the gene *GPBP1L1* that was selected in both analyses (2 and 3 views) was never reported in any cancer study.

### 3.3 Investigation on the selected features

#### 3.3.1 Differential gene expression, overrepresentation and gene set enrichment analyses

Differential gene expression analysis was performed between each pair of glioma types to get a more deep insight about the differences in gene expression between the classes. It was also performed to compare the results with the genes selected by DIABLO and understand whether this method gives additional and more useful information than the one that could be directly obtained by just a single-omics analysis. Figure 3A shows the volcano plots of the differentially expressed genes (DEGs) between each pair of glioma types, where the top 15 genes (with the lowest FDR) are labelled. Given the similarity between the LGG types, the DEGs obtained for astrocytoma *vs*. oligodendroglioma have a considerably lower log(FC) than the ones comparing the other classes. As expected, all the mRNA features selected by DIABLO were included in at least one of the sets of DEGs. Nevertheless, it is interesting to verify that not all the top DEGs were selected by DIABLO as the most relevant features to discriminate between the classes.

**Figure 3:**
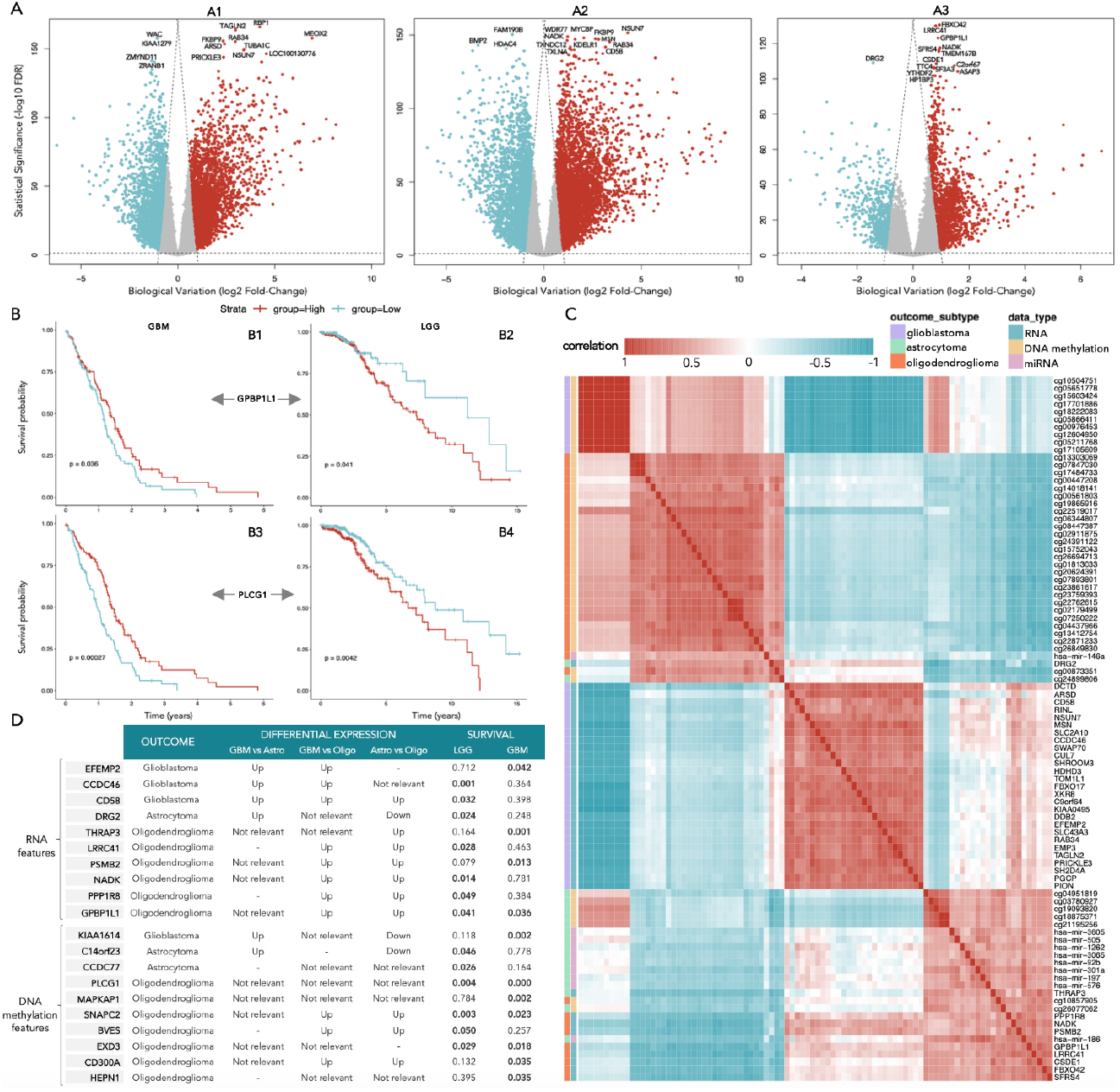
Differential gene expression, correlation and survival analysis with the selected features. A: Volcano plots showing the differential expressed genes, where the top 15 genes (with lowest FDR) are labelled, between: A1: GBM and astrocytoma; A2: GBM and oligodendroglioma; A3: Astrocytoma and oligodendroglioma. The red dots represent the positively expressed genes, while the blue dots represent the negatively expressed genes, in the type that is mentioned first relatively to the second. B: Survival analysis of the gene *GPBP1L1* in GBM (B1) and LGG patients (B2) and of the methylated gene *PLCG1* in GBM (B3) and LGG patients (B4). C: Correlation plot with all the selected features annotated with the respective outcome and belonging block of data. D: Selected features with effect on survival probability (p-value of log-rank test) in either LGG or GBM patients (statistical significant results for a 0.05 significance level are highlighted in bold).

Furthermore, overrepresentation (ORA) and gene set enrichment analysis (GSEA) were performed to have a more comprehensive view of the pathways and biological processes that are distinguished between classes (for more details, see Supplementary Material). The genes that are overexpressed in GBM when comparing with both LGG types are mainly associated with biological processes such as defense response, cell adhesion, extracellular matrix (ECM) structure organization and ECM-receptor interaction, while the genes that are underexpressed in GBM enrich for biological processes related with cell-cell signaling, synapse (including its organization, modulation of its transmission and vesicle cycle), regulation of transport, especially monoatomic ion transmembrane transport, and regulation of membrane potential. The biological processes that are differentially upregulated in GBM when comparing only to astrocytoma are mainly cytokine-mediated signalling and leukocyte migration, while with oligodendroglioma include the regulation of immune system process, cell population and leukocyte proliferation, cell migration, motility and secretion and response to cytokine. When comparing the LGG types, the upregulated genes in astrocytoma enrich for biological processes such as cytokine production, hemopoiesis, regulation of immune and defense response, cell and leukocyte proliferation and migration, while the downregulated genes are associated with cell-cell signaling, homeostasis, membrane potential and transport, synapse, nervous system and head development.

#### 3.3.2 Correlation analysis

In the previous subsections, the presence of several high correlations between features of different modalities was detected, when observing the two circos plots in Figures 1 and 2. Therefore, a correlation plot was further obtained with all the selected features, to understand not only the inter-block but also the intra-block correlations, as shown in Figure 3. There is a block that clearly stands out, with 10 methylation sites (between cg10504751 and cg17105609 (Figure 3C)) that are located respectively in the genes *GNAO1, MRC2, ARNTL, MCF2L2, TMEM106A, FGFRL1, KCNB1, KIAA1614, FES, FGFRL1*, positively correlated with each other and negatively correlated with the group of 27 genes (between *DCTD* and *PION* (Figure 3C)). All that features are associated with the distinction of GBM, and suggest involvement in common biological processes or cellular components. Some other smaller blocks with high positive correlations can be detected, such as the one composed by the methylation sites cg13303069, cg07847030 and cg17484733, all located in the gene *TCF7L1* and the one composed by the methylation sites cg22762615, cg02179499 and cg07250222, all located in the gene *FGFR2*. Both these blocks are negatively correlated with the group of genes *GPBP1L1, LRRC41, CSDE1, FBXO42* and *SFRS4*, which are associated with the oligodendroglioma type.

#### 3.3.3 Survival analysis

Additionally, to understand the possible prognosis value of each of the selected features, the Kaplan-Meier survival curve was then plotted using their gene expression levels. It was performed for GBM and LGG patients separately, given the remarkably different survival times from the TCGA cohorts: generally lower than 5 years for GBM but reaching 15 years in LGG patients, and to compare the different expression profiles in the two groups. The p-values of the log-rank test used to assess the statistical significance of the survival differences are summarized in Figure 3D, listing only the features which have a significant effect on survival in at least one of the groups. Notably, there are four different genes whose expression has an impact on survival probability in both groups of patients: *GPBP1L1, PLCG1, SNAPC2* and *EXD3*. Figure 3B exemplifies the Kaplan-Meier curves of *GPBP1L1*, selected within the mRNA features, and *PLCG1*, selected within the DNA methylation features (with p-value lower than 0.01 in both groups of patients). On the one hand, the higher expression of these genes reflects a higher survival probability in GBM patients (left side), while on the other hand, it has the opposite effect in LGG patients, by negatively impacting their survival probability. It is also interesting to note that some genes have a very significant effect (p-value <0.01) in the survival of one group of patients while its expression is helpful to distinguish another type. For instance, the expression of *CCDC46* gene impacts the survival of LGG patients, but is useful to discriminate the GBM type, while in the case of *THRAP3* gene, we have the reverse situation, distinguishing oligodendroglioma.

## 4 Discussion

Over the last years, the classification of glioma tumors has been subject of significant alterations as a result of the evolving understanding of its biology derived from extensive research into the molecular profiling of glioma types. The significant role of genetic markers is being increasingly highlighted in the determination of the final diagnosis, and the publicly accessible multi-omics databases, such as TCGA, are particularly helpful in gaining new knowledge regarding glioma characterization. However, the diagnostic categories reported for the TCGA-LGG and -GBM projects datasets are still not entirely consistent with the recently published WHO-2021 guidelines [7, 8]. In this study, we used the recently revised glioma reclassification and integrate multiple data modalities (including RNA-seq, DNA methylation, and miRNA) to gain more understanding of the biological mechanisms underlying glioma heterogeneity and discover novel biomarkers that can help further characterize each type and therefore be helpful for diagnosis and prognosis of the patients.

Using DIABLO, we were able to find predictive models that classify the three glioma types with high accuracy and identify relevant features in their characterization. This method showed to be more valuable than just applying a single-omics analysis like differential gene expression, where the correlation between features of different modalities is not considered. This network of connections between data layers has thus allowed to extract not only the most differentially expressed genes between classes, but the ones that are important for the separation of the three types, considering also the importance of DNA methylation and miRNA features and the relation established between each other.

It is known that GBM tumors are characterized by a highly malignant profile with increased cell proliferation and aggressiveness relatively to the LGG types, presenting significantly lower survival times even after chemotherapy and radiation treatments [40]. In this study, two highly correlated groups of molecular features were found to distinguish GBM from LGG, with the expression of the methylated sites positively correlated with each other and, in turn, negatively correlated with the selected genes. Interestingly, all these genes were revealed to be upregulated in GBM when compared with both astrocytoma or oligodendroglioma, while the methylation sites showed a considerably lower average level of methylation. It is worth noting that *KIAA0495*, which was selected with the highest loading in the distinction of GBM, codes a long non-coding RNA located at the arm of chromosome 1p, the absence of which is characteristic in oligodendroglioma. Nevertheless, the same gene is also downregulated in astrocytoma, where 1p deletion events are not so prevalent. In fact, *KIAA0495* has been suggested to modulate cellular apoptosis by regulation of p53-dependent antiapoptotic genes and to promote brain glioma proliferation and invasion, being associated with worse patients’ prognosis [41]. Other selected features including *XKR8, EMP3* and *GNAO1* also showed to be involved in the apoptosis process, but only the role of *EMP3* was already elucidated in glioma. The overexpression of *EMP3* in GBM is likely associated with the lack of *EMP3* hypermethylation that is present in LGG types [42], and its most thoroughly researched function is its regulation of receptor tyrosine kinase (RTK) signalling. The phosphorylation of tyrosine residues in signaling proteins is catalyzed by activated RTKs, starting multi-step signaling cascades which promote differentiation and proliferation. In particular, *EMP3* has been shown to foster the phosphorylation of the RTKs *EGFR* and ErbB2/HER2, as well as their downstream effectors (ERK, PI3K, Akt). Moreover, as a result of the *EGFR* gene amplification typically present in IDH-wt GBM, *EGFR* is frequently overactivated in this type, which thus correlates with increased proliferation, apoptosis resistance, and migration of tumor cells [43]. A set of other selected features, including *MSN, EFEMP2 GNAO1, TOM1L1, FES, FGFRL1* and *TMEM106A*, was also reported to participate in RTK signalling pathways. It is important to highlight the association of *CD58, CUL7, MRC2, SHROOM3* and the above-mentioned *FES, MSN, EFEMP2, GNAO1* in the extracellular matrix (ECM) organization, and, in special, the role of the last three in the integrin pathway. The ECM act as a biomechanical scaffold and a biochemical regulator of tumor cell homeostasis and is a crucial part of the GBM tumor microenvironment (TME). The ability of GBM tumor cells to migrate by inducing changes in actin cytoskeleton dynamics and ECM remodelling is well documented [44]. Integrins, as cell adhesion transmembrane receptors, act as ECM-cytoskeletal linkers that transmit signals between cells and the environment. Indeed, tumors can take advantage of the integrin-facilitated biological communication to take part in every stage of cancer progression, including tumor initiation, proliferation, and invasiveness [45]. In particular, while the *MSN* gene encodes an Ezrin-radixin-moesin (ERM) family protein that connects the actin cytoskeleton to the plasma membrane, *EFEMP2* was reported to be associated with the expression of matrix metalloproteinases (MMPs) [46], another group of proteins crucial in tissue remodelling. Although never described in glioma studies, *EFEMP2* and *GNAO1* are reported in pathway databases to be also involved with phospholipase-C (PLC) activity. PLC is stimulated by the P2Y2 receptor when coupled to specific G proteins, to hydrolyze PIP2, which is involved in calcium response, modulates a variety of actin binding proteins and activates Rac1 and RhoA [44]. These are small GTP-binding proteins of the Rho family known to act as molecular switches to control actin cytoskeleton dynamics. Interestingly, *GNAO1* is also linked to G-protein signaling and calcium regulation, and other selected features *SWAP70* and *MCF2L2* are players in Rho GTPase cycle. Furthermore, the role of *GNAO1* in purinergic signaling should be further investigated, since this pathway has been emerging as an important factor giving glioma cells invasive potential and resistance to ATP-induced cell death [47]. In this context, it has been proposed that besides alterations in the extracellular nucleotide/nucleoside metabolism, the disruption of purinergic signalling creates an inflammatory microenvironment with increase in cytokine production (specifically in IL-1*β*, IL-6 and TNF) and regulated platelet function [47]. Notably, several of the selected features are involved in either nucleotide metabolism or transport (*DCTD* and *SLC43A3*), in platelet function (*TAGLN2*) or in cytokine production or regulation (*CD58, TMEM106A, FES, MSN* and *GNAO1*). It is interesting to highlight that *TMEM106A*, a transmembrane protein found to be expressed on the surface of macrophages, specifically induces the release of IL-1*β*, IL-6 and TNF upon activation of the MAPK and NF-kappaB signaling pathways. This gene was never reported in glioma studies, but showed to be a tumor suppressor in gastric, renal and lung cancer [48], and therefore might also be also considered a novel marker candidate in glioma. Most of the remaining selected features are associated with metabolism, hemostasis and transport. For instance, *ARSD* that encodes the protein arylsulfatase D, essentially involved in sphingolipid, estrogen and protein metabolism, only gained attention recently for its role in amyloidosis [49]. Given that amyloid build-up is a part of the glioma tumor environment and it was inclusively indicated as a potential target for developing a novel class of anti-tumor drugs [50], more investigation should be dedicated to *ARSD* gene, whose function was never described in glioma.

Although all these genes were revealed to be relevant in biological processes underlying GBM and especially in its highly proliferative and invasive profile, the survival analysis showed that only the expression of the genes *CCDC46* (with updated symbol *CEP112*), *CD58, EFEMP2* and *KIAA1614* were significant in patient survival probability, the first two in LGG and the last two in GBM types. In fact, even though these genes are upregulated in GBM patients, they might not directly influence their survivability and still have an impact on LGG. Indeed, *CD58* was already reported before to be a prognostic biomarker in LGG [51]. It is worth noting that besides the involvement in the centrosome cycle, the function of *CCDC46* and *KIAA1614* in glioma is still unknown and deserves future research.

In contrast to GBM, LGG tumors typically show a more indolent course. However, many might eventually transform into a more aggressive type [52]. Further characterization of the different omics profiles can possibly offer new insights on tumor microenvironment and development from lower grade to higher grade gliomas. The current classification of LGG types is based mainly on two genetic markers, where the difference between astrocytoma and oligodendroglioma still relies only on the absence of 1p/19q codeletion [7]. In this study, multiple genetic features, including mRNA, DNA methylation and miRNA, were found to discriminate between astrocytoma and oligodendroglioma. Most of the selected genes were differentially expressed between GBM and oligodendroglioma but not between GBM and astrocytoma, with the exception of the gene *DRG2*, where the opposite was verified. In general, the expression of those genes was downregulated in oligodendroglioma when comparing with either astrocytoma or GBM, while the expression of *DRG2* is upregulated in both oligodendroglioma and GBM when comparing with astrocytoma. Also, there are just a few selected sites where the level of methylation presents a considerable difference between astrocytoma and both the other two types. Therefore, astrocytoma seems to be the more difficult type to define and separate, with large similarities with GBM and oligodendroglioma, while these two are more easily to distinguish. Nonetheless, it is still possible to highlight some few exceptional features that were selected *DRG2, KCND2* and *C14orf23*, which allow for the separation of astrocytoma. Indeed, *DRG2* was already reported before to be typically under-expressed in IDH-mutant samples, characteristic of LGG types, explaining its difference to GBM. This gene encodes a GTP-binding protein which catalyzes the conversion of GTP to GDP, and is known to be involved in the regulation of cell growth and differentiation. Although its function in glioma is still not completely elucidated, it was recently demonstrated to be positively correlated with several steps in anti-tumor immune response [53], and its depletion was shown to promote survival [54]. In the case of *KCND2*, it showed a lower level of methylation and a higher expression level in astrocytoma. It encodes a voltage-gated potassium channel that mediates transmembrane potassium transport in excitable membranes, primarily in the brain, regulating neurotransmitter release. Moreover, it also participates in signalling pathways such as ERK [55], important for cellular proliferation, and GDNF [56] (Glial cell line-derived neurotrophic factor), which guides glioma-associated microglia/macrophages (GAMs) recruiting in tumor immune resistance [57].

Furthermore, *C14orf23*, also known as *LINC01551*, is a lncRNA that influences cell cycle and transcription. It was never described in glioma, but it was suggested to promote metastatic ability by post-transcriptional regulation of miRNA (by ““sponge adsorption”) [58]. It would be interesting to further investigate the relationship between this gene and the miRNA selected as key players in glioma in order to understand its role in metastic progression.

From the selected features that distinguish oligodendroglioma, with down-regulated gene expression and higher level of methylation compared to the other types, one should note that the majority are related either with ubiquitination processes (*LRRC41, FBXO42, PSMB2*), RNA-binding activity, especially mRNA processing and splicing (*CSDE1, SFRS4, PPP1R8, GPBP1L1, THRAP3, SNAPC2, NANOS1*) or signalling (*LRRC41, PSMB2, PPP1R8, DRG2, GPR156, KCND2, BCL9L, PLCG1, MAPKAP1, FGFR2*). The signalling pathways influenced by these genes are mainly associated with RTK cascades, including the involvement of ERK, PI3K, Akt, EGFR and Rho GTPases, that were already discussed previously to be associated with increased proliferation and migration of tumor cells, and thus with a more invasive glioma profile. Additionally, other signalling pathways such as Notch and Wnt were also found to be essential in type distinction. For instance, *PSMB2* and *BCL9L* are known to be involved in the Wnt pathway, which activation was verified to increase the stemness of glioma cells [59]. On the other hand, *FBXO42* not only participates in ubiquitination and degradation of p53/TP53, but it was also shown as a critical regulator of the Notch pathway via modulation of RBPJ-dependent global chromatin landscape changes in leukemia [60]. It would be of great importance to further research the function of *FBXO42* in glioma, since it was only reported in other types of cancer. Interestingly, this gene is targeted by (hsa-)miR-186-5p, such as *LRRC41* and *CSDE1*. MicroRNAs (miRNAs) have recently attracted interest and their potential impact on oncogenic processes has been thoroughly studied. Based on altered miRNA expression profiles, it has been possible to identify and diagnose various tumors, as well as to forecast their development, prognosis, and response to treatment. For example, the selected miR-186 was already described in multiple cancers, including glioma, with involvement in the regulation of inflammatory response and apoptosis [61]. Among the other selected miRNA, it is also highlighted miR-92b, which also targets the selected gene *CSDE1* and *GPBP1L1*. This miRNA was indicated to restrain the proliferation, invasion, and stimulate apoptosis of glioma cells by targeting PTEN/Akt signaling pathway, suggesting a possible antitumor effect in glioma treatment [62].

Multiple genes revealed to have an important prognostic value, with significant effects on survival probability of LGG (including the genes *DRG2, LRRC41, NADK, PPP1R8* and *C14orf23*) and GBM patients (including the genes *THRAP3, PSMB2* and *MAPKAP1*). Notably, four genes showed to impact both LGG and GBM survival: *GPBP1L1, PLCG1, SNAPC2* and *EXD3. GPBP1L1, SNAPC2* and *EXD3* are likely key players in transcription, where the former is predicted to enable DNA and RNA binding and regulate transcription, the second encodes a subunit of the snRNA-activating protein complex, associated with the TATA box-binding protein, which is necessary for RNA polymerase II and III dependent small-nuclear RNA gene transcription, and the latter is involved in genetic stability and correction of DNA polymerase errors. While *SNAPC2* was already identified in glioma studies [63], *EXD3* was only reported in gastric cancer [64] and *GPBP1L1* was never reported in any cancer study. *PLCG1*, on the other hand, plays an important role in the intracellular transduction of receptor-mediated tyrosine kinase activators, already discussed to be fundamental in actin reorganization and cell migration. It is however very intriguing the relation of the expression profiles of *GPBP1L1* and *PLCG1* with the survival probability observed between GBM and LGG patients, in Figure 3B, with low expression affecting survival in GBM and high expression affecting survival in LGG. This suggests that the role of these genes and/or their interactions might be different in the two types, yet impacting patient survival in both cases. In fact, the expression of *PLCG1* was demonstrated to be significantly correlated with IDH status [65], which is one of the key characteristics that distinguish LGG and GBM.

## 5 Conclusions

In this study, we used the recently updated glioma reclassification to further characterize the different glioma types: GBM, astrocytoma and oligodendroglioma. Based on the integration of multiple omics’ layers (mRNA, DNA methylation and miRNA data from TCGA) using a supervised and sparse variant of canonical correlation analysis (DIABLO), we were able to discriminate the three glioma types with very high performance. Indeed, the correlation between blocks showed to be of extreme importance in the selection of the features. The group of correlated features that revealed to be relevant in the distinction of GBM from LGG types were mainly associated with RTK signalling and extracellular matrix organization, which is indeed a crucial part of the GBM tumor microenvironment. On the other hand, the discrimination of the LGG types was characterized mainly by features involved in ubiquitination and DNA transcription processes.

Furthermore, we were able to identify several novel features, that deserve future attention. For instance, the methylation of the gene *KIAA1614* is a potential biomarker with both diagnosis and prognosis value in GBM, yet its function is still not elucidated in glioma. Moreover, the gene *GPBP1L1* has a great impact in both LGG and GBM patient survival and showed distinct downregulation in oligodendroglioma when compared to both astrocytoma and GBM, but its role in cancer is still not described. Nonetheless, this gene is known to be targeted by miR-92b, a glioma-reported miRNA which was also selected by our method as a relevant feature to distinguish LGG types. Therefore, the interaction between the selected genes and miRNA features might also be essential in further investigation of these novel biomarkers.

## Supplementary information

The supplementary file contains the results from the Overrepresentation and Gene Set Enrichment Analyzes, for all pairs of glioma types using GO terms and KEGG pathways.

## Acknowledgements

The results presented here are based upon data generated by The Cancer Genome Atlas (TCGA) Research Network.

## Authors’ contributions

Francisca G. Vieira: Conceptualization, Methodology, Data analysis, Writing – original draft. Regina Bispo: Conceptualization, Methodology, Writing - review and editing, Supervision. Marta B. Lopes: Conceptualization, Methodology, Writing – review and editing, Supervision, Funding acquisition.

